# A single-cell transcriptomic analysis of the mouse hippocampus after voluntary exercise

**DOI:** 10.1101/2023.08.03.551761

**Authors:** Aditi Methi, Md Rezaul Islam, Lalit Kaurani, M Sadman Sakib, Dennis M. Krüger, Susanne Burkhardt, David Liebetanz, André Fischer

## Abstract

Exercise has been recognized as a beneficial factor for cognitive health, particularly in relation to the hippocampus, a vital brain region responsible for learning and memory. Previous research has demonstrated that exercise-mediated improvement of learning and memory in humans and rodents correlates with increased adult neurogenesis and processes related to enhanced synaptic plasticity. Nevertheless, the underlying molecular mechanisms are not fully understood. With the aim to further elucidate these mechanisms we provide a comprehensive dataset of the mouse hippocampal transcriptome at the single-cell level after four weeks of voluntary wheel-running. Our analysis provides a number of interesting observations. For example, the results suggest that exercise affects adult neurogenesis by accelerating the maturation of a subpopulation of *Prdm16*-expressing neurons. Moreover, we uncover the existence of an intricate crosstalk among multiple vital signaling pathways such as NF-κB, Wnt/β-catenin, Notch, retinoic acid (RA) pathways altered upon exercise in a specific cluster of excitatory neurons within the Cornu Ammonis (CA) region of the hippocampus. In conclusion, our study provides an important resource dataset and sheds further light on the molecular changes induced by exercise in the hippocampus. These findings have implications for developing targeted interventions aimed at optimizing cognitive health and preventing age-related cognitive decline.

## Introduction

The positive effects of exercise in maintaining both physical and cognitive health have been widely established during the past few decades. In addition to alleviating the risk of metabolic diseases (S. A. Phillips et al., 2015; Sylow & Richter, 2019; Zanuso et al., 2010), cardiovascular disease (Gielen et al., 2015; Morris, 1994; S. A. Phillips et al., 2015) and several types of cancers (Galvão & Newton, 2005; Hojman et al., 2018), the benefits of physical exercise on brain function have also been extensively studied and reviewed (Fischer, 2016; Hillman et al., 2008; Praag, 2009; Sutoo & Akiyama, 2003; Voss et al., 2011). Numerous studies in model organisms have shown that exercise promotes significant structural and functional changes in the hippocampus, which is a region of the brain that is essential for learning and memory processes. These changes indicate an overall positive impact of exercise on hippocampal function, including protective effects against aging and neurological disorders (Ashdown-Franks et al., 2020; Brown et al., 2013; Cass, 2017; Fischer et al., 2007; Mahalakshmi et al., 2020; Sujkowski et al., 2022; Swenson et al., 2020; Zhao et al., 2020), increased adult neurogenesis (Pereira et al., 2007; Praag, Christie, et al., 1999; Praag et al., 2005; Praag, Kempermann, et al., 1999; Seri et al., 2001), enhanced synaptic plasticity and improved cognitive performance (Cotman & Berchtold, 2002; Erickson & Kramer, 2009). Multiple studies have also demonstrated these beneficial effects in humans, indicating that aerobic exercise leads to an increase in the size of the human hippocampus, as well as improvements in memory performance (Cassilhas, 2012; Erickson, 2011; Maass, 2015; Voss, 2013).

While the molecular mechanisms underlying these effects are not fully understood, exercise was shown to increase the production of several neurotrophic factors (Fabel et al., 2003; Loprinzi & Frith, 2019; Neeper et al., 1995; Trejo et al., 2001) and other signaling molecules, such as neurotransmitters and cytokines (Lin & Kuo, 2013; Małkiewicz et al., 2019; Meeusen & Meirleir, 1995; Meeusen & Piacentini, 2001; Packer et al., 2010; Pedersen, 2000). Exercise was also found to induce changes in gene expression measured via quantitative polymerase chain reaction (qPCR), microarray or RNA sequencing. For example, exercise has been found to upregulate the expression of genes that are involved in cell proliferation and differentiation, as well as genes that are involved in enhancing neuronal and synaptic plasticity (Cotman & Berchtold, 2002; Mojtahedi et al., 2013; Tong et al., 2001). Many of these studies have focused on rodent models and have used running wheels as a behavioral paradigm to test the effects of physical exercise (Pereira et al., 2007; Praag, Christie, et al., 1999; Praag et al., 2005; Praag, Kempermann, et al., 1999; Rendeiro & Rhodes, 2018).

Recent studies have demonstrated that single nuclei RNA sequencing (snRNA-seq), which is a powerful next generation sequencing (NGS) technique and has several advantages over traditional RNA sequencing methods (such as enabling transcriptomic analysis of tissues that are difficult to dissociate or have low cell yields), enables detection of previously unknown cell types and transcriptional subtypes (Gouwens, 2020; Habib, 2017; Lake, 2018; Mathys, 2021; Yao, 2021). Furthermore, it can be leveraged to better understand adult neurogenesis at the single cell level in both mice and humans (Habib, 2017; Zhou et al., 2022). Although a number of previous studies have looked into the transcriptomic and epigenetic changes induced by exercise on the brain (Benito et al., 2018; Buckley et al., 2022; Goldberg et al., 2021; McGreevy et al., 2019; Miguel et al., 2021; Pan-Vazquez et al., 2015; Shah et al., 2017; Urdinguio et al., 2021; Y. Y. Wang et al., 2023), to our best knowledge, none have investigated hippocampal cell-type specific transcriptomic changes following four weeks of voluntary exercise in mice without employing additional behavioral tests. In this study, we used running wheels as a form of voluntary exercise and aimed to further elucidate exercise-related mechanisms in the hippocampus using snRNA-seq.

Our results indicate that exercise selectively affects the proportions of cells in specific clusters of neuronal cell-types and suggest that the transcription factor *Prdm16* may play a key role in orchestrating the maturation of newborn dentate gyrus neurons upon exercise. In addition, our data hint to an important role crosstalk between NF-κB, Wnt/β-catenin, Notch and retinoic acid (RA) signaling pathways in a specific excitatory neuron cluster belonging to the Cornu Ammonis (CA) region, which could collectively regulate and enhance synaptic plasticity in these neurons upon exercise. In conclusion, our findings provide an important novel dataset and help to further elucidate the molecular processes associated with improved neuronal plasticity in response to aerobic exercise.

## Results

### snRNA-seq analysis of the mouse hippocampus reveals changes in cell-type abundance upon exercise

To understand how physical exercise affects the hippocampus, we designed an experiment where 8 wild type (WT) mice were divided into 2 equally sized groups - runners that had free and voluntary access to running wheels in their cages (the ‘exercise’ group), and sedentary mice that had similar wheels in their cages that were however blocked (the ‘control’ group) **(Supplementary Table 1)**. This voluntary exercise paradigm lasted 4 weeks, after which we isolated whole hippocampal nuclei from all mice for snRNA-seq analysis **(Fig. 1A)**. A total of 22817 nuclei across the 8 samples were recovered after quality control, doublet removal and filtering (see methods). After accounting for batch effects, clustering was performed and nuclei were grouped into 21 unique clusters. Subsequently, these clusters were assigned to major hippocampal cell types including excitatory neurons (ExN), inhibitory neurons (InN) as well as microglia, astrocytes, oligodendrocytes and oligodendrocyte precursor cells based on the expression of cell-type specific marker genes **(Fig. 1B, C, Supplementary Fig. 1 and Supplementary Table 2)**. Excitatory neurons made up the majority of cells, followed by oligodendrocytes and inhibitory neurons **(Fig. 1D)**. Additionally, we also identified excitatory neuron clusters that belonged to either the CA regions (ExN2-10 and ExN12-13) or the dentate gyrus (ExN1 and ExN11) within the hippocampus, using marker genes for these regions (Cembrowski et al., 2016) **(Fig. 1C, E and Supplementary Table 2)**. Additionally, distinct marker genes for each of these clusters were determined computationally (see methods) **(Supplementary Table 3)** and gene ontology (GO) analyses of the top markers specific to each of the neuronal clusters suggested different biological functions for these neurons within the hippocampus **(Supplementary Fig. 2 and Supplementary Tables 4-6)**.

**Fig. 1:**
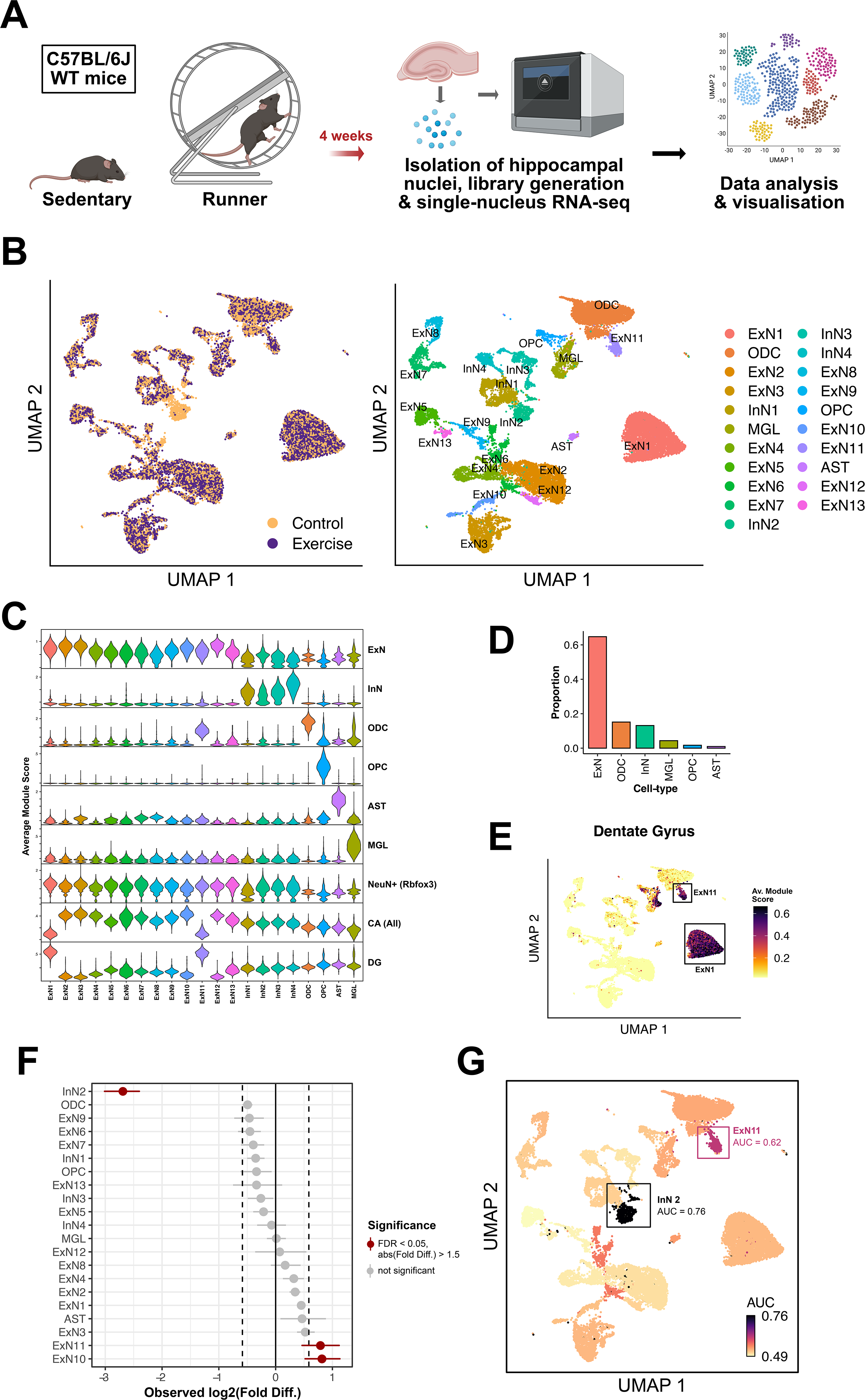
Single-nucleus RNA sequencing analysis of the whole hippocampus in exercising vs. sedentary mice reveals changes in abundance of specific cell-types. **A.** Experimental design: The cohort of WT mice belonging to the exercising experimental group, or ‘runners’ (*n* = 4), had free access to running wheels in their cages for a duration of 4 weeks. Mice belonging to the control or ‘sedentary’ group (*n*lJ=lJ4) were similarly housed but the running wheels were blocked. The 4 week-long voluntary exercise paradigm was followed by isolation of hippocampal nuclei for single-nucleus RNA sequencing. **B.** UMAP plots showing clusters of nuclei or ‘cells’, colored by experimental group (left panel) and cell-type (right panel). *(ExN1-13: excitatory neurons; InN1-4: inhibitory neurons; ODC: oligodendrocytes; OPC: oligodendrocyte precursor cells; AST: astrocytes; MGL: microglia)* **C.** Violin plots showing average module score (expression) of marker genes specific to the different cell types/hippocampal regions, after cell-type specific annotation of clusters *(ExN: excitatory neurons; InN: inhibitory neurons; ODC: oligodendrocytes; OPC: oligodendrocyte precursor cells; AST: astrocytes; MGL: microglia; CA: Cornu Ammonis; DG: Dentate Gyrus).* **D.** Bar graph indicating the proportions of broad cell-types observed in the dataset *(ExN: excitatory neurons; InN: inhibitory neurons; ODC: oligodendrocytes; OPC: oligodendrocyte precursor cells; AST: astrocytes; MGL: microglia)* **E.** UMAP plot with cell clusters colored by average module score (expression) of marker genes specific to the dentate gyrus (DG), highlighting the two DG excitatory neuron clusters. **F.** Analysis of differences in cell-type proportions between cells from exercise and control samples, using permutation testing. The x-axis denotes the fold difference in cell-type proportions (log2 scale). Points marked in red indicate clusters with significantly different proportions of cells between the two groups. Horizontal lines around the points indicate the confidence interval for the magnitude of difference for a specific cluster, calculated via bootstrapping. **G.** UMAP plot with cells colored by Augur cell-type prioritization upon perturbation resulting from exercise, measured using area under the curve (AUC) scores.

Next, we applied permutation testing to analyze if exercise would lead to detectable differences in proportions of cells originating in each cluster. Out of all clusters, InN2, ExN10 and ExN11 showed significant changes in cell-type abundance (**Fig. 1F**). Using Augur (Squair et al., 2021), we also performed cell-type prioritization analysis to detect which cell types are most perturbed in response to exercise. To measure the perturbation levels, we employed the area under the receiver operating characteristic curve (AUC). An AUC value of 0.5 indicates that cells from the exercise condition in a cluster have no significant difference in perturbation compared to cells from the control condition (random chance). Conversely, a value of 1.0 signifies that every cell from the exercise condition exhibits higher perturbation compared to the control condition. Augur performs subsampling in a way that changes in relative abundances of cell-types across conditions do not confound the analysis. Hence, if certain cell-types are found to be strongly perturbed due to exercise, it could imply that differences in cell-type proportions appear due to these perturbations and not the other way around. We observed that clusters InN2 and ExN11 were the only two clusters with an AUC value > 0.6, indicating changes in cell-type proportions within these clusters upon exercising (**Fig. 1G**). More specifically, upon exercise fewer cells were detected for cluster InN2 while more cells were detected within cluster ExN11. In particular, the increased proportion of cells within cluster ExN11 may reflect increased adult neurogenesis, which has been repeatedly described upon exercise (Pereira et al., 2007; Praag, Christie, et al., 1999; Praag et al., 2005; Praag, Kempermann, et al., 1999; Seri et al., 2001). Thus, while the changes observed for cluster ExN11 may indicate the increased number of excitatory neurons upon neurogenesis, the changes seen within cluster InN2 are more difficult to explain. Therefore, we decided to first analyze cluster InN2 in greater detail.

### Exercise induces changes in an inhibitory neuron population expressing *Prdm16*

We identified four distinct inhibitory neuron clusters in the dataset based on the overall expression of marker genes Gad1 and Gad2. Each cluster was furthermore characterized by high specific expression of the following genes: Sox6 (InN1), Prdm16 (InN2), Cnr1 (InN3), and Egfr (InN4) **(Supplementary Fig. 3)**. As indicated in Fig. 1E, we observed a selective reduction in abundance (log2 fold difference = −2.69) of inhibitory neurons in cluster InN2 in the exercise condition, as compared to the controls (**Fig. 2A**). Further functional analysis of the top marker genes for InN2 cells distinguishing them from the other inhibitory clusters (i.e. differentially expressed (upregulated) in InN2 as compared to InN1, InN3 and InN4) broadly implicated genes involved in transmembrane transport of ions as well as dendritic spine development (**Fig. 2B and Supplementary Table 7**). Among the top 20 of these markers, a few were more distinctly expressed in InN2 as compared to the rest. These included *Prdm16*, *Ano1*, *Ano2*, *Zfhx3, Zic1*, *Zic4*, *Zfp521* and *Ankfn1* (**Supplementary Fig. 4A**). *Prdm16* is a transcription factor that belongs to the PRDM family of transcriptional regulators. It has been widely characterized as an important cell-fate switch regulator in brown adipose tissue (Kajimura et al., 2010; Seale et al., 2008, 2011), but recent studies have also identified its role in neural stem cell homeostasis (including maintenance of the neural stem cell pool), as well as development, differentiation and positional specification of neurons (Aguilo et al., 2011; Chuikov et al., 2010; Leszczyński et al., 2020; Shimada et al., 2017; L. Su et al., 2020; Turrero García et al., 2020). *Ano1* and *Ano2* are calcium-activated chloride channels (CaCCs) of the anoctamin protein family. CaCCs play crucial roles in regulating the excitability of smooth muscle cells and also some types of neurons (Hawn et al., 2021). Specifically, *Ano1* has been shown to contribute to the process maturation of radial glial cells during cortex development (Hong et al., 2019), while *Ano2* modulates action potential waveforms in hippocampal neurons, along with the integration of excitatory synaptic potentials (Huang et al., 2012). Moreover, *Ano2* has been implicated in calcium-dependent regulation of synaptic weight in GABAergic inhibition in the cerebellum, hence regulating ionic plasticity (Zhang et al., 2015). *Zfhx3* is a large transcription factor with 23 zinc fingers and 4 homeodomains (Morinaga et al., 1991), which has been shown to promote neuronal differentiation during neurogenesis in development and primary cultures (Jung et al., 2005; Miura et al., 1995). *Zic1* and *Zic4* are also transcription factors containing zinc finger domains, which are involved in nervous system development (Gaston-Massuet et al., 2005; Inoue et al., 2007), specifically by regulating neuron differentiation and maintaining neural precursor cells in an undifferentiated state (Inoue et al., 2007). *Zfp521* encodes another zinc-finger protein that regulates many genes involved in neural differentiation. Studies have shown that *Zfp521* is essential and sufficient for driving the intrinsic neural differentiation of mouse embryonic stem cells (Kamiya et al., 2011), and is also sufficient for converting adult mouse brain-derived astrocytes into induced neural stem cells (Zarei-Kheirabadi et al., 2019). Little is known about *Ankfn1* but it has been genetically linked to cannabis dependence (Agrawal et al., 2011).

**Fig. 2:**
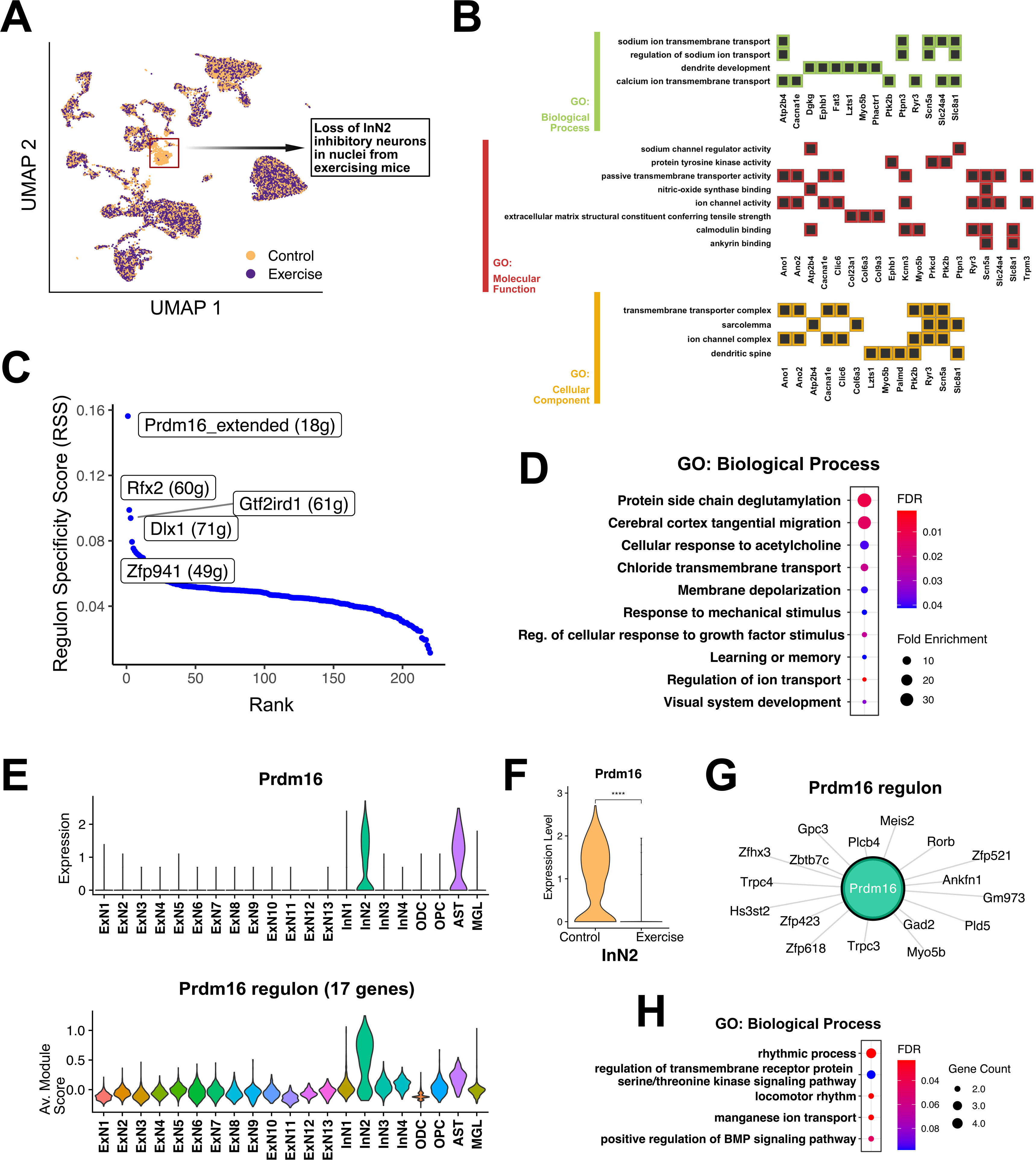
Selective loss of InN2 inhibitory neuron cluster upon exercise indicates a role of the transcription factor Prdm16. **A.** UMAP plot with cells colored by experimental group, showing the loss of InN2 inhibitory neurons upon exercise. **B.** Heatmap plots of functional annotation for specific genes among the top 50 markers of the InN2 cluster that had significantly enriched GO (biological process, molecular function and cellular component) terms. **C.** Cell-type specific regulons for InN2 cluster identified using the SCENIC workflow. The y-axis denotes the regulon specificity score (RSS) (with high RSS values indicating high cell-type/cluster specificity, and vice versa). The x-axis denotes the rank of each regulon within the selected cluster, based on the RSS. The top 5 ranked regulons for InN2 are labeled on the plot, with the number of genes comprising each regulon indicated within parentheses. *(Regulons ending with “extended” also include motifs linked to the transcription factor by lower confidence annotations)*. **D.** Dotplot showing significant GO biological process terms enriched among the collective list of genes and transcription factors (TFs) making up the top 5 regulons, as indicated in the RSS-Rank plot in (C). **E.** Violin plots depicting the normalized expression of Prdm16 **(top panel)**, and the average module score (expression) for the genes included in the Prdm16 regulon **(bottom panel)** in all clusters. **F.** Violin plot showing the expression of Prdm16 in InN2 cluster, split between exercise and control cells. **G.** Network plot showing the 17 genes comprising the Prdm16 regulon. **H.** Dotplot showing significant GO biological process terms enriched among the list of 17 Prdm16 regulon genes.

In order to further understand what makes the InN2 inhibitory neurons distinct, we used the SCENIC workflow (Aibar et al., 2017) to identify regulons (sets of transcription factors and their putative targets) that were specifically active in the InN2 cluster (**Fig. 2C and Supplementary Table 8**). A functional enrichment analysis of the collective list of genes comprising the top 5 InN2-specific regulons (*Prdm16*, *Rfx2*, *Gtf2ird1*, *Dlx1* and *Zfp941*) indicated significant enrichment of gene ontology (GO) terms for regulation of synaptic signaling and transmembrane ion transport, membrane depolarization and response to acetylcholine, and interestingly, regulation of learning and memory (**Fig. 2D and Supplementary Table 9**). Among these top 5 regulons, the transcription factor *Prdm16* and its 17 target genes comprised the most specific active regulon in this cluster. This was substantiated by the highly specific expression in InN2 of both *Prdm16* and the combined expression of the 17 regulon genes (**Fig. 2E**). The observed reduction of exercise cells in InN2 was subsequently indicated in the stark upregulation of Prdm16 in the control cells (**Fig. 2F**). 5 candidates from the *Prdm16* regulon (*Prdm16*, *Ankfn1*, *Meis2*, *Myo5b*, *Zfhx3*, *Zfp521*) are also shared with the top 20 InN2 marker genes (differentially expressed in InN2 as compared to InN1, InN3 and InN4) (**Supplementary Fig. 4**), most of which have been implicated in regulating neuronal differentiation (Agoston et al., 2014; Jung et al., 2005; Kamiya et al., 2011; Y. Liu et al., 2013; Miura et al., 1995; L. Su et al., 2020; Turrero García et al., 2020). Additionally, functional analysis specifically on the *Prdm16* regulon genes suggests enrichment of GO terms such as rhythmic process (*Ankfn1*, *Hs3st2*, *Rorb*, *Zfhx3*) and regulation of bone morphogenetic protein (BMP) signaling (*Prdm16*, *Gpc3*, *Zfp423*), which is known to modulate neuronal maturation and adult neurogenesis (Bond et al., 2012) (**Fig. 2G, 2H and Supplementary Table 10**).

Among the other top regulons, the gene candidates within the *Dlx1* regulon indicated that *Dlx1* itself positively regulates *Prdm16*. *Dlx1* is a TF that, along with its related gene *Dlx2*, is involved in regulating neuronal migration (of newborn neurons, away from the proliferative zone) as well as the functional longevity of GABAergic interneurons in the hippocampus (Cobos et al., 2005). Moreover, a recent study indicated that the expression of the *Meis2* gene driven by *Dlx1/2* promoted the fate determination of striatal neurons in mice (Z. Su et al., 2022). Since *Meis2* is regulated by *Prdm16* (as seen in the *Prdm16* regulon), and it is also one of the top markers of the InN2 cluster which is comprised mostly of control cells, these findings collectively suggest that exercise leads to repression of *Prdm16* in specific hippocampal neurons (which are characterized by high *Prdm16* expression). Although we identified these neurons initially as cluster InN2, it is possible that these cells represent a specific developmental stage during adult neurogenesis that is only captured in control mice in our dataset, since exercise is known to increase proliferation but also maturation of adult newborn neurons (Ambrogini et al., 2013; Lattanzi et al., 2022; Wu et al., 2008; Yang et al., 2015). In conclusion, these data may suggest that decreased expression of *Prdm16* could be an important regulatory step during adult neurogenesis that is affected by exercise.

### Exercise affects a subpopulation of excitatory neurons linked to enhanced adult neurogenesis

Following the cell-type abundance and perturbation analyses (Fig. 1E and F), we also sought to take a deeper look into the ExN11 cluster belonging to the dentate gyrus. These cells showed an increase in abundance (log2 fold difference = 0.79) in the exercise condition, as compared to the controls. This increase can also be seen in a small fraction of cells from ExN11 that seem to integrate into the mature granule cell cluster ExN1 of the dentate gyrus (as indicated in Fig. 3D by expression of GC markers *Prox1* (Cembrowski et al., 2016; Erwin et al., 2020; Kempermann et al., 2015) and *Calb1* (Kempermann et al., 2004), which shows an overall low but relatively higher expression in ExN1), with the number of these integrating cells increasing more than two-fold after exercise (**Fig. 3A**).

**Fig. 3:**
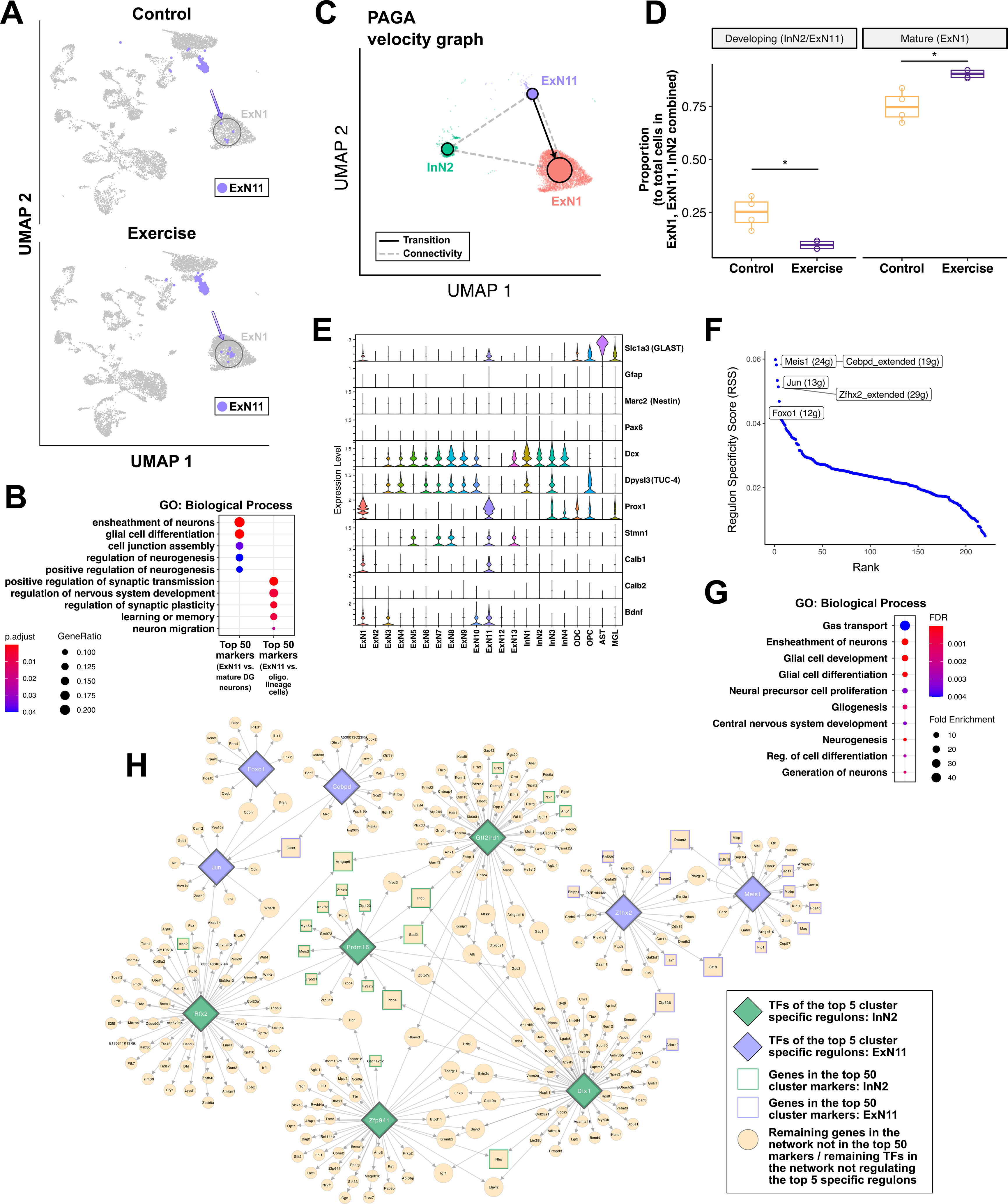
Increased abundance of neurons in ExN11 cluster suggests increased neurogenesis upon exercise. UMAP plots highlighting the ExN11 cluster, split by the experimental conditions (exercise and control). **B.** Dotplot showing significant GO biological process terms enriched among the top 50 genes differentially expressed between ExN11 and ExN1 clusters, and among the top 50 genes differentially expressed between the ExN11 cluster and the ODC and OPC clusters. **C.** Partition-based graph abstraction (PAGA) graph for InN2, ExN11 and ExN1 clusters, with velocity directed edges constructed from RNA velocity measurements. Edges denote either connectivities (dashed) or transitions (solid/arrows). **D.** Box-plots showing a significant decrease in the proportion of developing cells (ExN11/InN2) and an increase in the proportion of mature granule cells (ExN1) upon exercise. Each dot indicates one sample (n=4/4 for exercise and control samples). **E**. Violin plots showing normalized expression of selected marker genes for radial glia/immature neurons/exercise-mediated neurogenesis. **F.** Cell-type specific regulons for ExN11 cluster identified using the SCENIC workflow. The y-axis denotes the regulon specificity score (RSS) (with high RSS values indicating high cell-type/cluster specificity, and vice versa). The x-axis denotes the rank of each regulon within the selected cluster, based on the RSS. The top 5 ranked regulons for ExN11 are labeled on the plot, with the number of genes comprising each regulon indicated within parentheses. **G.** Dotplot showing significant GO biological process terms enriched among the collective list of genes and transcription factors (TFs) making up the top 5 regulons, as indicated in the RSS-Rank plot above. **H.** Combined gene-regulatory network plot with TFs specific to InN2 and ExN11 clusters, and their respective regulon genes (Larger nodes represent genes/TFs that connect two or more regulon networks).

In order to understand the underlying gene expression patterns of ExN11 cells, we looked at the top computationally detected marker genes for this cluster. Most of these markers were not specific to ExN11, and in fact were also highly expressed in either ExN1 neurons, or oligodendrocyte lineage cells (ODC and OPC clusters). This becomes more evident after plotting the combined expression of the top 50 genes differentially expressed between ExN11 and ExN1 (**Supplementary Fig. 5A and Supplementary Table 11**), and the top 50 genes differentially expressed between ExN11 and the oligodendrocyte lineage clusters (ODC and OPC) (**Supplementary Fig. 5B and Supplementary Table 12**). Functional enrichment analysis of these two lists of differentially expressed markers indicates enrichment of GO terms related to ensheathment of neurons, regulation of synaptic plasticity, neurogenesis and neuronal migration (**Fig. 3B and Supplementary Table 13**), suggesting that increased neurogenesis upon exercise results in more excitatory neurons in ExN11.

Next, we generated a partition-based graph abstraction (PAGA) graph with velocity directed edges constructed from RNA velocity measurements (Bergen et al., 2020; Wolf et al., 2019), after subsetting the dataset to analyze our clusters of interest: InN2, ExN11 and ExN1 (**Fig. 3C**). The graph indicates a potential connectivity between these three clusters, and a transition from ExN11 to ExN1, which supports our findings so far suggesting that ExN11 represents newborn excitatory neurons that eventually integrate into the dentate gyrus. Moreover, we see that the proportion of cells coming from ExN11 and InN2 combined, taken among the total cells from the three clusters of interest (ExN1, ExN11 and InN2), decreases in exercise, while the proportion of mature granule cells (ExN1) increases in the exercise condition as compared to controls (**Fig. 3D**). This suggests that exercise leads to faster maturation of developing or immature neurons (InN2 and ExN11) to mature granule cells (ExN1).

We also looked at the expression of markers corresponding to different stages of neurogenesis in this cluster, in order to assign these cells to a specific stage of adult neurogenesis (Bohlen Und Halbach, 2007) (**Fig. 3E**). However, we observed either no expression (*Neurod*, *Tbr2*, *Tubb3*) or very low expression (*Gfap*, *Nestin*, *Pax6*) of many canonical markers, possibly due to dropout effects in the sequencing data. As compared to other neurons, ExN11 cells show higher expression of *GLAST* (*Slc1a3*), which is a marker for radial glia cells, but at the same time we also observed low expression of Dcx and TUC-4 (*Dpysl3*) and high expression of *Prox1* which could suggest that these are transiently amplifying (or transit amplifying) progenitor cells (Potten & Loeffler, 1990; Torii et al., 1999). This latter cell-type is known to show transient expression of markers: initially expressing glial or stem cell markers like *Gfap* and *Nestin*/*Sox2*, and later expressing granule cell markers such as *Prox1*. ExN11 cells also express stathmin (*Stmn1*) at relatively higher levels. Stathmin is a microtubule destabilizing protein that is considered to be an immature neuron marker, since it has been implicated in controlling the transition from dividing neuronal precursors to postmitotic neurons in the subgranular zone of the DG during adult neurogenesis (Boekhoorn et al., 2014). *Bdnf (brain-derived neurotrophic factor)*, which is known to be involved in exercise-mediated neurogenesis in the hippocampus (Isackson et al., 1991; P. Z. Liu & Nusslock, 2018; Neeper et al., 1995; Sleiman et al., 2016; Vaynman et al., 2004), showed very low expression throughout the dataset, which made it difficult to reliably quantify any overall changes in its expression levels in the exercise condition as compared to the sedentary controls. However, it did have relatively higher expression levels in ExN11, although we found no significant differences between the exercise and control conditions in this cluster.

We further analyzed the gene regulatory landscape for ExN11 using the SCENIC workflow to identify regulons which are specifically active in this cluster, and found the *Meis1* regulon to be the most specific out of the top 5 regulons (**Fig. 3F and Supplementary Table 14**). *Meis1* is a transcription factor that has been shown to play a role in promoting neuronal differentiation and possibly also neurogenesis (Barber et al., 2013; Mojsin & Stevanovic, 2009; Owa et al., 2018). Other top transcription factors include *Cebpd (CCAAT enhancer binding protein delta)* which has been implicated in hippocampal neurogenesis and regulation of learning and memory (Banerjee et al., 2019; Powell et al., 2017), *Jun* which is an immediate early gene and transcriptional regulator involved in cell proliferation (K. H. Schlingensiepen et al., 1994; K.-H. Schlingensiepen et al., 1993; Weston & Davis, 2007), *Zfhx2* which is involved in neuronal differentiation (Komine et al., 2012), and *Foxo1* which regulates the long-term maintenance of adult neural stem cells, and belongs to the FoxO family that plays an important role in the maintenance of autophagic flux and neuronal morphogenesis in adult neurogenesis (Ro et al., 2013; Schäffner et al., 2018). A functional enrichment analysis of the collective list of genes comprising these top 5 ExN11-specific regulons indicated significant enrichment of gene ontology (GO) terms for gas transport (*Car2*, *Car14* - carbonic anhydrases that play a role in neuronal excitability and signaling (Ruusuvuori & Kaila, 2014)), regulation of neural development, neurogenesis, and also glial cell differentiation, which is in line with our previous observation of glial markers in this cluster (**Fig. 3G and Supplementary Table 15**). We also observed that some of the TFs from the top regulons of ExN11 and InN2 clusters shared downstream target genes, and also regulated each other, such as *Rfx2* regulating *Jun* as well as *Prdm16*. These common gene-regulatory relationships between our two clusters of interest are highlighted in **Fig. 3H**. Building on our previous hypothesis, these results further suggest that the observed loss of *Prdm16*-expressing neurons upon exercise might represent a molecular snap shot of enhanced adult neurogenesis and especially the maturation of new born neurons. In this scenario, ExN11 neurons could be a consequence of their differentiation into and possible switch to an excitatory neuron subtype.

### Differential gene expression and gene regulatory network analyses reveal broad transcriptional changes linked to synaptic plasticity

To get a better understanding of the broad transcriptomic changes mediated by exercise, we looked at differential gene expression between cells from the exercise and control conditions in all other clusters apart from InN2 and ExN12 (complete list of differentially expressed genes is given in **Supplementary Table 16**). Most clusters revealed either no significant changes in gene expression or very few deregulated genes across the two conditions, with the exception of ExN1, ExN2, ExN6, InN1 and ODC (**Fig. 4A** (top) and **4B** (top)). The top significant GO (biological process) terms enriched for these deregulated genes indicate broad processes involved in specific clusters upon exercise (**Fig. 4A** (bottom) and **4B** (bottom), **Supplementary Tables 17-18**). In particular, ExN6 shows a distinctly large number of both up- and downregulated genes, with the former being enriched for dendrite morphogenesis and synaptic signaling pathways and the latter for postsynapse organization and synapse assembly processes (**Supplementary Tables 19-20**). UpSet plots in Fig. 4A and 4B identified certain genes that are commonly up- or downregulated upon exercise across multiple clusters. *Vps13a*, for example, is upregulated in exercise in 8 out of the 21 clusters and also downregulated in 2 excitatory neuron clusters, suggesting a role of chorein (the protein encoded by *Vps13a*) which is a powerful regulator of cytoskeletal architecture and cell survival in many cell types (Föller et al., 2012). Moreover, *Vps13a* mutations lead to chorea-acanthocytosis, a rare disorder that also affects the brain and reduced chorein levels have been linked to Alzheimer’s disease (Lazarczyk et al., 2016). *Plxna4*, which has been shown to regulate synaptic plasticity (Tan et al., 2019) and dendrite morphogenesis in the hippocampus (Yamashita et al., 2016), is upregulated in ExN1 and ExN6. *Zbtb16*, upregulated in oligodendrocytes and ExN7, is involved in neural progenitor cell proliferation and neuronal differentiation during development (Usui et al., 2021). *Fkbp5* is a glucocorticoid receptor (GR) binding protein, which negatively regulates GR, and is interestingly upregulated upon exercise only in oligodendrocytes and microglia. It has been associated with mental disorders as well as in regulating neuronal synaptic plasticity (Qiu et al., 2019). Among the downregulated genes, *Ddx5* is an RNA helicase which has been shown to regulate neurogenesis and neuronal differentiation, likely by inhibiting reprogramming to pluripotency (H. Li et al., 2017; Suthapot et al., 2022). *Tcf4*, downregulated in ExN1 and InN1, has been linked to neurodevelopment (Forrest et al., 2014), regulation of dendritic spine morphology (Crux et al., 2018) and regulation of neuronal migration through *Bmp7* (Chen et al., 2016). Altered *Tcf4* function has also been linked to schizophrenia and mice in which *Tcf4* is overexpressed develop cognitive impairment, which is attenuated by exercise (Badowska et al., 2020).

**Fig. 4:**
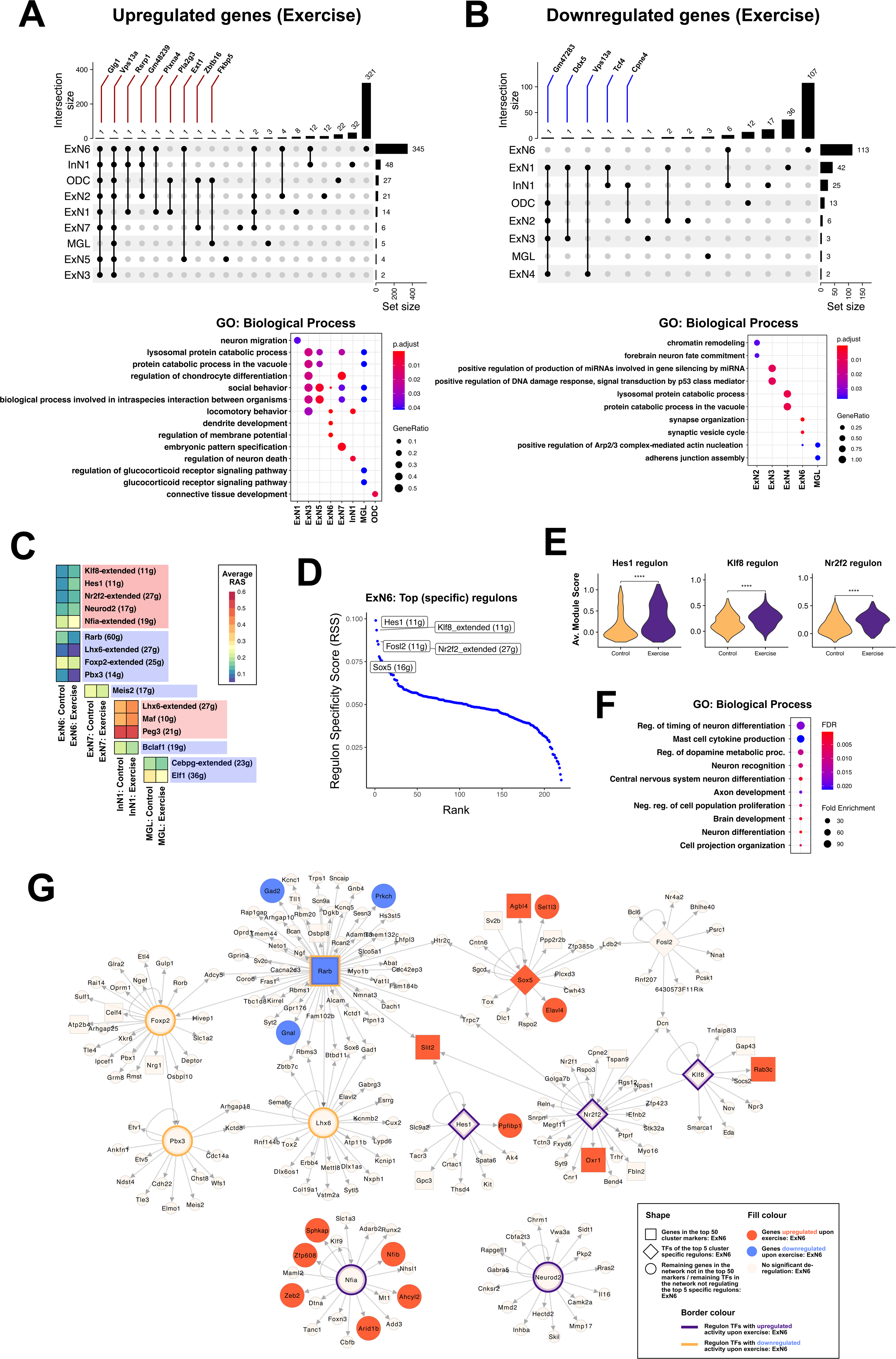
Differentially expressed genes in excitatory neurons suggest regulation of synaptic plasticity and neuron differentiation to promote increased neurogenesis upon exercise. **A. (top panel)** and **B. (top panel)** UpSet plots depicting (A) commonly upregulated and (B) commonly downregulated genes upon exercise, between different clusters. Each row corresponds to a cluster, and bar charts on the right show the size of the set of genes up-/downregulated upon exercise in that cluster. Each column corresponds to a possible intersection (commonly up-/downregulated gene(s)): the filled-in cells show which cluster set is part of an intersection. Gene names are labeled for intersections with < 2 common genes among clusters. **A. (bottom panel)** and **B. (bottom panel)** Dotplots showing GO biological process terms enriched among (A) genes significantly upregulated and (B) genes significantly downregulated upon exercise, in clusters with significantly enriched terms. **C.** Heatmap depicting the average regulon activity scores (RAS) for regulons with significant differential activity between cells from exercise and control conditions, in specific clusters. The number of genes comprising each regulon is indicated within parentheses. Regulons within red boxes indicate those upregulated upon exercise, and the ones within blue boxes indicate the ones downregulated upon exercise. **D.** Cell-type specific regulons for ExN6 cluster identified using the SCENIC workflow. The y-axis denotes the regulon specificity score (RSS) (with high RSS values indicating high cell-type/cluster specificity, and vice versa). The x-axis denotes the rank of each regulon within the selected cluster, based on the RSS. The top 5 ranked regulons for ExN6 are labeled on the plot, with the number of genes comprising each regulon indicated within parentheses. **E.** Violin plots showing the average module score (expression) of the 3 regulons specific to ExN6 that also show upregulated activity upon exercise (*Hes1*, *Klf8*, *Nr2f2*). **F.** Dotplot showing significant GO biological process terms enriched among the collective list of genes and transcription factors (TFs) making up the top 5 regulons, as indicated in the RSS-Rank plot in (C). **G.** Network plot with TFs and their respective regulon genes, specific to or showing differential activity in ExN6 upon exercise, indicating gene regulatory interactions in ExN6 cluster (Genes/TFs that are significantly deregulated upon exercise or are specifically overexpressed in ExN6 are represented bigger in size, while selected genes belonging to these regulons which are deregulated in the same direction as the regulon TF are colored in red or blue).

We further identified regulons that significantly differ in their activity across the exercise and control cells in certain clusters (**Fig. 4C**). Functional enrichment analysis of these regulon genes suggests that these TFs regulate a variety of biological processes in selected cells upon exercise (**Supplementary Fig. 6A and Supplementary Table 21**). Since ExN6 clearly exhibits a larger number of regulated genes as well as regulons with differential activity upon exercise, we sought to characterize these neurons by looking at the specific transcription factors active in this cluster, using the SCENIC workflow. The top 5 ExN6-specific regulons included *Hes1*, *Klf8*, *Nr2f2*, *Fosl2* and *Sox5* (**Fig. 4D and Supplementary Table 22**), and the top 3 regulons (*Hes1*, *Klf8* and *Nr2f2* regulons) also showed significant upregulation upon exercise in the differential regulon activity analysis as shown in Fig. 4C, further corroborated by increased overall expression of these regulon genes in the ExN6 exercise cells (**Fig. 4E and Supplementary Fig. 6B**). The functional enrichment analysis of the combined list of genes comprising the top 5 regulons suggested regulation of timing of neuron differentiation (mediated by *Hes1* and *Sox5*) as the term with the highest enrichment (**Fig. 4F and Supplementary Table 23**). Additionally, *Hes1* is an important effector gene of the Notch signaling pathway, which has been linked to learning and memory by regulating synaptic plasticity in mature neurons (Costa et al., 2003; Y. Wang et al., 2004). Interestingly, *Hes1* was also induced in the early (0-1h) time window upon acute exercise in skeletal muscle tissue (Amar et al., 2021), and another study showed that inactivation of *Hes1* in excitatory neurons resulted in abnormal fear and anxiety behaviors (Matsuzaki et al., 2019). *Klf8* is a TF playing critical roles in various biological processes such as regulation of cell cycle progression. Elevated protein levels of *Klf8* were shown to induce the activation of Wnt/β-catenin signaling pathway, by promoting the expression and stabilization of β-catenin which further interacts with and inhibits NF-κB to regulate synaptic plasticity and cognition (Yi et al., 2014). Moreover, Wnt/β-catenin signaling has been strongly implicated in the regulation of synaptic assembly, neuroprotection, neurotransmission as well as synaptic plasticity, while disruption or deregulation of Wnt signaling has been linked to several neurodegenerative diseases such as AD (Oliva et al., 2013; Yi et al., 2014). *Nr2f2* (or COUP-TFII) belongs to the COUP-TF transcription factor family which is involved in regulating neuronal differentiation and migration of neural progenitors (Fuentealba et al., 2010), and was additionally observed to be a member of the *Klf8* regulon from the ExN6 SCENIC analysis. Interestingly, the regulon for retinoic acid (RA) receptor *Rarb* showed decreased activity in exercise cells, along with the regulons for *Lhx6*, *Foxp2* and *Pbx3* (as seen in Fig. 4C). *Rarb* is a direct transcriptional target of RA signaling, and this suggests its possible role in regulating neuronal plasticity upon exercise, which could be linked to the already-established roles of RA signaling in neuronal differentiation (Janesick et al., 2015).

These findings suggest a pattern of crosstalk between NF-κB, Wnt/β-catenin, Notch, RA and other signaling pathways, which could collectively regulate and enhance synaptic plasticity specifically in ExN6 neurons upon exercise. These relationships are visualized and highlighted in the ExN6 gene regulatory network (**Fig. 4G**).

## Discussion

In this study, we aimed to investigate the effects of physical exercise, in the form of voluntary wheel running, on the mouse hippocampal transcriptome at the single-cell level. Voluntary wheel running is an established approach in rodent models to test the effects of aerobic exercise on the brain (Pereira et al., 2007; Praag, Christie, et al., 1999; Praag et al., 2005; Praag, Kempermann, et al., 1999; Rendeiro & Rhodes, 2018). We decided to investigate the hippocampal transcriptomic changes at the 4 week time point after exercise, since this experimental duration has commonly been employed in similar studies and has been shown to be sufficient in enhancing neuronal plasticity and would moreover allow us to capture specific states of adult neurogenesis (Kodali et al., 2016; Miguel et al., 2021; Pan-Vazquez et al., 2015).

Through our analysis, we first observed that 4 weeks of exercise training leads to changes in the proportions of specific cell-types, most significant and pronounced in two clusters: a reduction in cells in the inhibitory cluster InN2 and an increase in cells in the excitatory cluster ExN11. Further investigation into the gene expression patterns in InN2 indicated that these cells, almost all originating from the control (sedentary) samples, showed high expression of the TF *Prdm16*. *Prdm16* has already been studied in the context of cell-fate switch regulation in brown adipose tissue, but interestingly, many recent studies have implicated its potential role in neural development and differentiation, particularly in stem cell homeostasis and maintenance and positional specification of neurons (Aguilo et al., 2011; Chuikov et al., 2010; Leszczyński et al., 2020; Shimada et al., 2017; L. Su et al., 2020; Turrero García et al., 2020). For example, one study showed that *Prdm16* in mouse medial ganglionic eminence (MGE) progenitors plays an essential role in controlling the timing of maturation of forebrain GABAergic interneurons, with *Prdm16* expression promoting the proliferation of interneurons and repressing maturation (Turrero García et al., 2020). Expression of other top gene markers for InN2 indicated genes such as CaCCs *Ano1* and *Ano2* regulating ion transport, neuronal excitability and plasticity, and zinc-finger protein encoding genes *Zfhx3*, *Zic1*, *Zic4* and *Zfp521* involved in neuronal and stem-cell differentiation. We further looked into the transcriptional regulation of InN2 by identifying highly active regulons in this cluster, with the results again implicating *Prdm16* and its target genes, which showed enrichment of GO terms involved in neuronal maturation and adult neurogenesis, such as rhythmic process (*Ankfn1*, *Hs3st2*, *Rorb*, *Zfhx3*) and regulation of BMP signaling (*Prdm16*, *Gpc3*, *Zfp423*). This was substantiated by the highly specific expression of the *Prdm16* gene in InN2, suggesting that the observed loss of exercise cells in InN2 is linked to the high expression of *Prdm16* in cells from the control samples. Moreover, *Dlx1*, another highly active InN2 regulon that plays a role in regulating the functional longevity of GABAergic interneurons in the hippocampus, regulates *Prdm16* and drives the expression of *Prdm16*-regulated Meis2 gene. Interestingly, a recent study also indicated that high *Prdm16* expression is essential in promoting the disappearance of radial glia and the ending of cortical neurogenesis in the postnatal mouse brain (J. Li et al., 2023). Collectively, these findings, supported by the existing literature, allowed us to hypothesize that InN2 cells characterized by high *Prdm16* expression could represent a specific developmental stage during adult neurogenesis captured only in cells from sedentary mice, possibly as a result of faster maturation of these *Prdm16*-high neurons upon exercise. These data also suggest that snRNA-seq analysis upon exercise could be a suitable approach to detect distinct molecular stages of adult neurogenesis.

This view is further supported when we investigated the NeuN-positive cluster ExN11 that showed an increase in the number of cells after exercise. Through RNA velocity measurements and PAGA graph generation, we observed a potential cell-type transition (in the context of trajectory inference) from the ExN11 cluster to the mature DG cluster ExN1, suggesting that ExN11 represents newborn excitatory neurons that eventually integrate with the mature granule cells in the DG. ExN11 also seemed to express both neuronal and glial gene markers, which is consistent with studies that characterized radial glia-like precursor cells with astrocytic as well as neural stem cell-like properties during neurogenesis (Kempermann et al., 2015; Steiner et al., 2004). At the same time, ExN11 cells were *Dcx*-negative and had high expression of *Prox1*, suggesting that these could be transit amplifying progenitor cells that show transient expression of markers: from glial/stem cell markers to granule cell markers, while high stathmin (*Stmn1*) expression in these cells could also implicate them as postmitotic immature neurons (Boekhoorn et al., 2014). Upon exploring the gene regulatory landscape for ExN11, we further identified specifically active regulons such as *Meis1*, *Cebpd*, *Jun*, *Zfhx2* and *Foxo1*, all of whom are involved in neurogenesis, neuronal as well as glial differentiation, morphogenesis and development. It was interesting to note some common gene-regulatory relationships between ExN11 and InN2 clusters, such as *Rfx2* regulating *Jun* as well as *Prdm16*.

On this basis, we suggest that ExN11 neurons indicate increased adult neurogenesis upon exercise. Although the relevance of adult neurogenesis in humans is a topic that remains under debate and is still associated with inconsistencies, with some studies claiming that there was no evidence of neurogenesis past youth and in adulthood (Sorrells et al., 2018), there is also much evidence supporting adult hippocampal neurogenesis in humans (Eriksson et al., 1998) including studies implicating a decline of the latter with age (Boldrini et al., 2018; Spalding et al., 2013; Vecchio et al., 2018). Moreover, research has indicated that adult-born hippocampal neural stem cells can be induced to produce distinct neuronal populations, and these neurons, despite losing their proliferative capacity and showing age-related inactivity, may be reactivated through certain mechanistic stimuli such as running (Katsimpardi et al., 2014; Katsimpardi & Lledo, 2018; Katsimpardi & Rubin, 2015; Lugert et al., 2010; Villeda et al., 2014).

Additionally, we propose that the observed loss of InN2 neurons and higher number of ExN11 neurons could potentially result from an inhibitory to excitatory switch of *Prdm16*-expressing developing GABAergic neurons after exercising. It has already been shown that the polarity switch of GABAergic responses from depolarization to hyperpolarization could be a physiological hallmark of neuronal maturation and that this dual role of GABAergic transmission is closely linked to neuronal development (Ben-Ari et al., 1989; Kim et al., 2012; Mueller et al., 1984; Owens & Kriegstein, 2002; Pozzi et al., 2020). At the same time, these developmental changes imply the existence of structural and functional changes in GABAergic transmission in the adult brain as compared to neonatal development. Therefore, even though the inferences from our results suggest a polarity switch in the opposite direction (inhibitory to excitatory), we believe this provides new insights into the involvement of GABA in neuronal differentiation and maturation in the context of exercise-induced neurogenesis and changes in plasticity in the adult hippocampus.

In summary, these data may indicate that downregulation of *Prdm16* in maturing new born hippocampal neurons could be an important regulatory step by which exercise enhances the process of adult neurogenesis. It could therefore be interesting to test strategies that reduce *Prdm16* function in the hippocampus as a therapeutic approach to improve cognitive function.

Finally, we also looked at the broad gene expression patterns in the dataset that differed between exercise and sedentary conditions in specific cell-types. Although most clusters only showed very mild gene expression changes between the two conditions, ExN6 (belonging to CA excitatory neurons) exhibited a markedly high number of regulated genes as well as regulons with differential activity upon exercise, prompting us to explore it further. The most differentially active as well as specific regulon in ExN6 was *Hes1*, an important effector gene of the Notch signaling pathway, which itself regulates learning and memory by regulating synaptic plasticity in mature neurons (Costa et al., 2003; Y. Wang et al., 2004). GO term analysis indicated a role of *Hes1* in regulating the timing of neuron differentiation, mediated along with another specific ExN6 regulon *Sox5*. Multiple studies have shown that oscillating *Hes1* expression regulates the maintenance and proliferative capacity of neuronal progenitor cells, acting as a repressor of the commitment of the latter to a neuronal fate, which in turn influences the number of neurons produced, the timing of neurogenesis as well as overall brain morphogenesis (Nakamura et al., 2000; Ochi et al., 2020; N. E. Phillips et al., 2016; Shimojo et al., 2011; Sueda & Kageyama, 2020). Interestingly, *Hes1* was also induced in the early (0-1h) window upon acute exercise in skeletal muscle tissue (Amar et al., 2021), further indicating the possibility of an oscillating pattern of Notch signaling activation upon exercise. Our findings suggest that *Hes1* upregulation (and subsequent Notch activation) in ExN6 at the 4-week exercise time point could mark a late maturation state of adult neurogenesis in adjacent DG neurons, potentially through lateral inhibition of neuronal fate commitment of newborn cells. This is in line with our previous hypothesis that *Prdm16* upregulation in InN2 neurons at this time could also be associated with an important maturation step in adult neurogenesis in the DG, and that ExN11 neurons represent a maturing state of excitatory cells that are past the neurogenesis phase. Analysis of other top gene markers and regulons of ExN6 (*Klf8*, *Nr2f2*, *Fosl2* and *Sox5*) suggested involvement of pathways regulating neuronal and synaptic plasticity, such as Notch signaling, retinoic acid (RA) signaling and Wnt/β-catenin signaling, which is further linked to inhibition of NF-κB to regulate synaptic plasticity and cognition. Together, these findings point to a unique pattern of crosstalk between these various signaling pathways, suggesting that they act collectively to regulate and enhance synaptic plasticity in ExN6 neurons upon exercise..

This study of course comes with its own limitations, such as technical drop-outs in the snRNA-seq dataset that limit gene expression analyses. We also look solely at a fixed snapshot of changes occurring at the 4-week time-point after exercise, and it would be beneficial to consider a longer range and/or multiple time-points for temporal changes that occur with these experimental paradigms, especially when it comes to gene expression, which would give a better insight into the dynamic processes we highlight with our study. Additionally, the different neuronal clusters/subtypes and their interactions could be more deeply characterized through functional studies in the future. Despite the fact that our findings are currently limited to interpretations from bioinformatic analyses and a single snRNA-seq dataset, we believe that they encourage further research focusing on functional interpretation and validation of these results, and probe important questions in the field of aerobic exercise and its impact on brain health.

In conclusion, our study provides a useful single-cell sequencing resource for exploring exercise-induced transcriptomic changes in the adult mouse hippocampus. Some of the observed changes allow for a deeper insight into the molecular processes that control adult hippocampal neurogenesis. For example, we provide evidence that a cluster of neurons characterized by high *Prdm16* expression may represent a specific state in the maturation of adult born hippocampal neurons and our data suggest that a decrease of *Prdm16* levels is critical to eventually promote the maturation of newborn neurons in the adult brain. We also identified a cluster of excitatory CA neurons exhibiting pronounced gene expression changes leading to regulation and enhancement of synaptic plasticity in the hippocampus upon exercise. Since adult hippocampal neurogenesis is impaired during aging and neurodegenerative diseases such as Alzheimer’s diseases, targeting *Prdm16* could be a novel approach to enhance plasticity in the diseased brain.

## Methods

### Animal cohort

The experiments were performed according to the ethical guidelines of the national animal protection law and were approved by the responsible ethical committee of the State of Lower Saxony, Germany. Male C57BL/6J mice were purchased from Janvier and housed individually in an animal facility with a 12-h light–dark cycle at constant temperature (23 °C) with *ad libitum* access to food and water. Animals were subjected to voluntary running at 3 months of age. Mice were allowed free access to the running wheel 24 h/day (Klinker et al., 2017). In the control groups, the running wheel was blocked.

### Sample preparation, single nuclei isolation, library preparation and sequencing

For RNA sequencing analysis 4 runners and 4 control mice at the age of 3 months were assessed. Sample preparation, isolation of nuclei and library preparation for sequencing were performed according to previously published protocols (Kettwig et al., 2021). After 4 weeks in cages with or without running wheels, mice were deeply anesthetized with ketamine and medetomidine and transcardially perfused with phosphate-buffered saline (PBS, pH 7.4). After cervical dislocation, the whole brain was removed from the skull and the preparation of the hippocampus was performed in a petri dish filled with ice cold PBS buffer under a binocular. Tissue samples from two hemispheres of different mice from the same behavioral cohort were pooled and immediately frozen on liquid nitrogen and stored at −80 °C until further processing. Nuclei from frozen mouse brain tissues were isolated according to the previously published protocol with certain modifications (Sakib et al., 2021). Briefly, frozen tissues were Dounce homogenized in 500 µl EZ prep lysis buffer (Sigma NUC101) supplemented with 1:200 RNAse inhibitor (Promega, N2615) for 45 times in a 1.5 mL Eppendorf tube using micro pestles. The volume was increased with lysis buffer up to 2000 µl and incubated on ice for 5 min. Lysates were centrifuged for 5 min at 500 × *g* at 4 °C and supernatants were discarded. The pellet was resuspended into 2000 µl lysis buffer and incubated for 5 min on ice. After 5 min of centrifugation (500 × *g* at 4 °C), the resulting pellet was resuspended into 1500 µl nuclei suspension buffer (NSB, 0.5 % BSA, 1:100 Roche protease inhibitor, 1:200 RNAse inhibitor in 1×PBS) and centrifuged again for 5 min (500 × *g* at 4 °C). The pellet was finally resuspended into 500 µl NSB and stained with 7AAD (Invitrogen, Cat: 00-6993-50). Single nuclei were sorted using BD FACS Aria III sorter. Sorted nuclei were counted in Countess II FL Automated Counter. Approximately 4000 single nuclei per sample were used for GEM generation, barcoding, and cDNA libraries according to 10X Chromium Next GEM Single Cell 3 Reagent v3.1 protocol. Pooled libraries were sequenced in Illumina NextSeq 550 in order to achieve >50,000 reads/nuclei.

### Preprocessing and exploratory analysis of snRNA sequencing data

The raw sequencing data generated from the snRNA-seq libraries were demultiplexed with cellranger mkfastq using CellRanger software v.4.0.0 (10XGenomics). Reads were aligned to the mm10 genome (NCBI:GCA_000001635.8) (GRCm38.p6) to obtain gene counts. The cellranger count pipeline was used with default options to generate a gene-count matrix by mapping reads to the pre-mRNA reference to account for unspliced nuclear transcripts.

All following downstream analyses were performed R (version 4.1.0). The feature-barcode matrices generated for all 8 samples by CellRanger were converted to sample-wise Seurat objects using the Run10X() and CreateSeuratObject() functions from the Seurat package (version 4.0.2) (Hao, 2021). For each sample, the object was filtered to remove nuclei where less than 200 genes and/or 500 reads and/or greater than 50000 reads were detected. Genes that were expressed in fewer than 10 nuclei were filtered out. Moreover, any nuclei expressing more than 0.1% mitochondrial gene counts (‘percent.mt’) were also removed. Normalization was done using the SCTransform() function from the Seurat and sctransform (version 0.3.2) packages (Hafemeister & Satija, 2019), with the arguments method = “glmGamPoi” and vars.to.regress = “percent.mt”. PCA embeddings were calculated using the RunPCA() function, and RunHarmony() function from the harmony package (version 0.1.0) (Korsunsky et al., 2019) was used to remove confounding effects due to sequencing batches. Leiden clustering (Traag et al., 2019) was performed using the FindNeighbors() and FindClusters() functions in Seurat with 50 principal components and a resolution parameter of 0.2. Uniform Manifold Approximation and Projection (UMAP) (McInnes et al., 2018) was used for visualization of clusters with the following arguments as parameters: umap.method = “umap-learn”, metric = “correlation”, n.neighbors = 30, n.components = 2, min.dist = 0.3, spread = 0.5, reduction = “harmony”, dims = 1:50. For each sample object, the DoubletFinder package (version 2.0.3) (McGinnis et al., 2019) was used to detect and remove any potential doublets. All individual sample objects were then merged into one Seurat object, which was re-normalized, integrated and re-clustered using the SCTransform, RunPCA(), RunHarmony(), FindNeighbors() and FindClusters() functions, with the same parameter values as before. UMAP was again used for visualization of clusters in the final merged object. Cell-type annotation of clusters was done using the expression of selected marker genes (as listed in **Supplementary Table 1**). Three small clusters (22, 23 and 24) identified after the initial clustering (**Supplementary Fig. 1A**) were removed from the dataset before further analysis, since they did not show specific expression of any cell-type markers and subsequently could not be assigned to any known cell-type.

### Cell-type abundance and prioritization analyses

The differences in the proportion of exercise and control cells in each cluster were calculated using the scProportionTest R package (https://github.com/rpolicastro/scProportionTest) (version 0.0.0.9), which calculates the p-value of the magnitude difference of abundance for each cluster using a permutation test (no. of permutations = 1000), and generates a confidence interval for the same using bootstrapping. Clusters with an observed fold difference magnitude of at least 1.5 with an FDR threshold of 0.05 were considered to have significant changes in cell-type proportions between exercise and control conditions.

To identify clusters (cell-types) in which the response to perturbation because of the given stimulus, i.e. exercise, is prioritized, we used the Augur R package (version 1.0.3) (Skinnider et al., 2021; Squair et al., 2021). This tool trains a machine-learning classifier specific to each cluster to predict whether the cells originate from exercise or control samples, based on the hypothesis that the separability between cells “responding” to exercise and the ones that don’t can be quantified by seeing how readily the experimental sample labels already associated with each cell (exercise or control) might be predicted from their underlying molecular measurements. The accuracy of the classifier is further subjected to cross-validation to strengthen the predicted results.

### Differential gene expression and regulon activity analyses

The top gene markers for each cluster, as well as genes differentially expressed between exercise and control cells in each cluster, were identified using the FindMarkers() function in Seurat with the following parameters: logfc.threshold = 0.1, test.use = “MAST”, latent.vars = c(“batch”,“percent.mt”), assay = “RNA”. We adopted the single-cell regulatory network inference and clustering (SCENIC) workflow (Aibar et al., 2017; Sande et al., 2020) to construct gene regulatory networks from our data and evaluate the activity of these gene regulatory networks in cells from different cell-type clusters, using the SCENIC R package (version 1.3.1). Briefly, this analysis was performed as follows: with the gene expression matrix as input, gene sets that were co-expressed with potential regulators or transcription factors (TFs) were identified using the GENIE3 algorithm (implemented using the GENIE3 R package (version 1.16.0) (Huynh-Thu et al., 2010)). Using the RcisTarget R package (version 1.14.0) (Aibar et al., 2017), *cis*-regulatory motif analysis was performed to further exclude false positives and indirect targets of TFs and get direct-binding targets. With this step, only those co-expression modules that showed significant motif enrichment of a potential regulator (TF) were selected, and these final modules of TFs and their putative downstream target genes were referred to as ‘regulons’. Finally, the AUCell algorithm (Aibar et al., 2017) was used to quantify the activity of these regulons in cells by assigning an enrichment or activity score to each regulon. This was calculated as an ‘area under the recovery curve’ (AUC) across the ranking (based on expression values) of all genes expressed in a cell. The top 5 regulons specifically active in a selected cluster (compared to other clusters) were identified using the calcRSS() and plotRSS_oneSet() functions in the AUCell R package (version 1.16.0). The differentially active regulons across exercise and control cells within a cluster were determined using the FindMarkers() function in Seurat by setting the argument assay = ‘AUC’, where the ‘AUC’ assay slot in the Seurat object contained the regulon AUC score matrix generated by AUCell.

For functional annotation and enrichment analysis of selected differentially expressed genes or differentially active regulons, an overrepresentation analysis was carried out using one of the following methods: clusterProfiler R package (version 4.0.0) (Yu et al., 2012), WebGestalt R package (version 0.4.4) (Liao et al., 2019) and the ShinyGO web application (Ge et al., 2020). Gene regulatory relationship networks were constructed using the output of the SCENIC workflow and the Cytoscape software (version 3.9.1) (Shannon et al., 2003).

### RNA velocity analysis and PAGA velocity graph generation

To understand cellular dynamics at a deeper level, we used the scVelo Python toolkit (version 0.2.4) (Bergen et al., 2020) with default parameters on our dataset, which utilizes splicing kinetics to calculate RNA velocity estimates for single cells. Using these velocity estimates, we used the PAGA graph abstraction method (Wolf et al., 2019) within the scVelo workflow to infer connectivities and directed transitions between selected cell-type clusters in our data. A PAGA graph was then visualized by associating a node with each cluster of interest and connecting these nodes based on the statistical measure of connectivity or transition between them.

## Data availability

snRNA-seq data is available via GEO accession number GSE234278.

## Acknowledgments

This work was supported by the following grants to A.F.: The DFG (*Deutsche Forschungsgemeinschaft*) SFB1286, Germany’s Excellence Strategy - EXC 2067/1 390729940 and the JPND project EPI-3E.

## Contributions

A.M., M.R.I. and A.F. conceptualized and planned the project and analysis. M.R.I. and D.L. organized and performed the behavioral experiments. M.R.I and M.S.S. prepared the samples for snRNA-seq and M.S.S. performed the nuclei isolation and sorting. L.K. and S.B. prepared the sequencing libraries and performed the snRNA sequencing. D.M.K. provided bioinformatics support and performed demultiplexing, alignment and count matrix-generation from the raw snRNA-seq data. A.M. analyzed the snRNA-seq data, visualized and interpreted results. A.M. and A.F wrote the manuscript, with input from other co-authors.

