## Supplementary Figures for "A single-cell transcriptomic analysis of the mouse hippocampus after voluntary exercise"

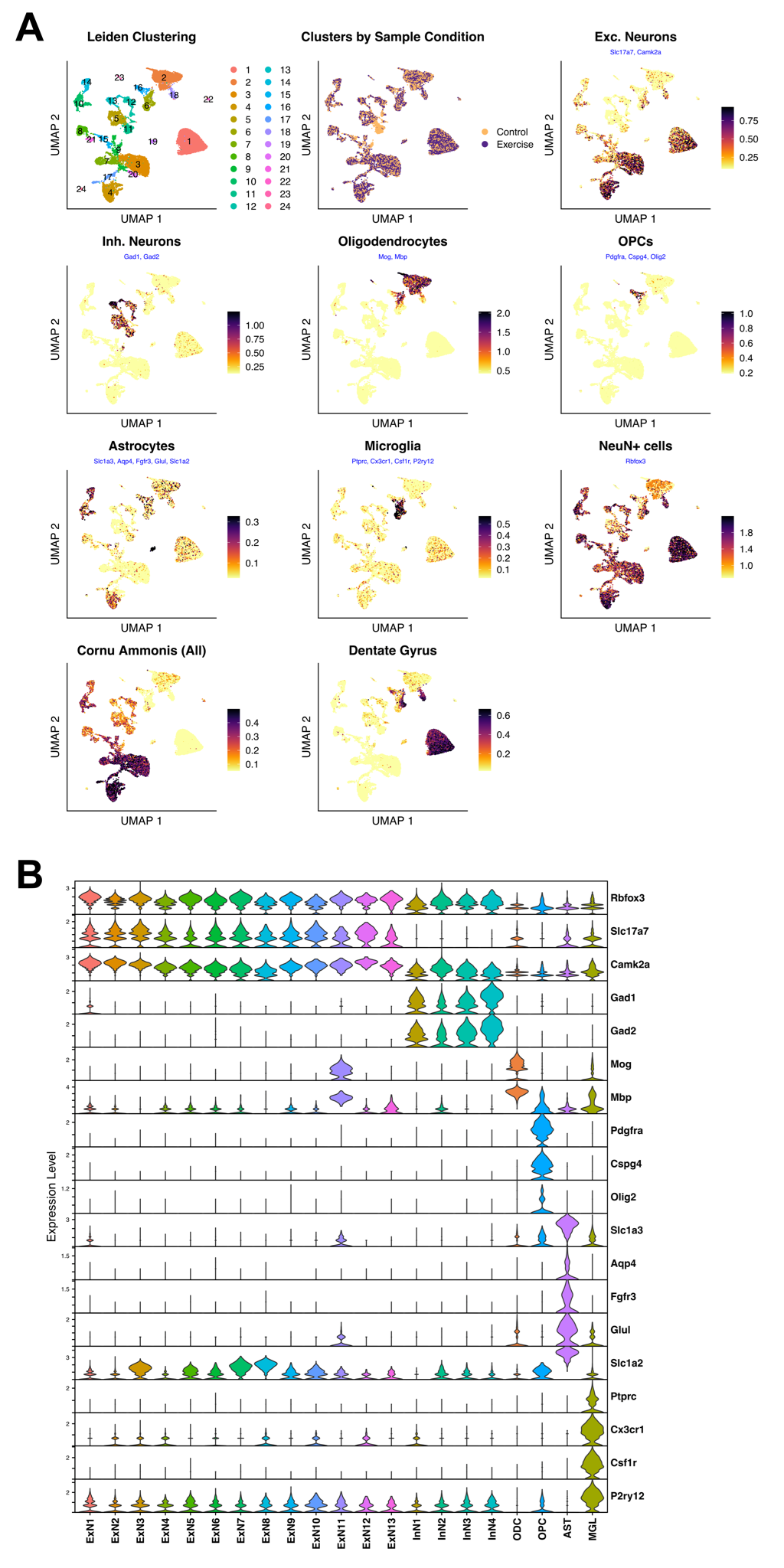

**Supp. Fig. 1: Cluster annotation. A.** UMAP plots with originally identified (unannotated) cell clusters colored by average module score (expression) of marker genes specific to the different cell types/hippocampal regions. **B.** Violin plots showing normalized expression of individual marker genes specific to the different cell types, after cell-type specific annotation of clusters *(ExN1-13: excitatory neurons; InN1-4: inhibitory neurons; ODC: oligodendrocytes; OPC: oligodendrocyte precursor cells; AST: astrocytes; MGL: microglia).*

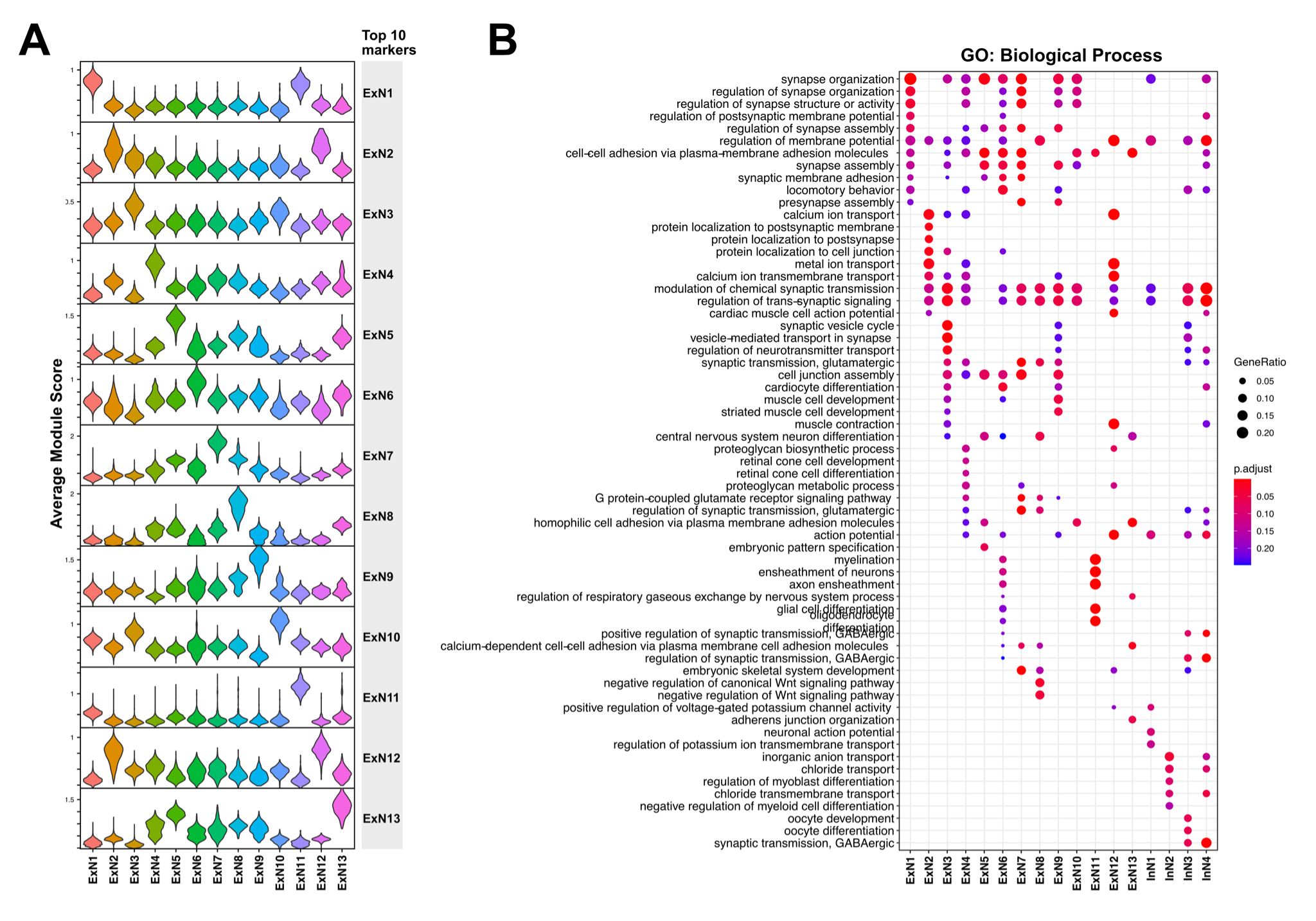

**Supp. Fig. 2: A.** Violin plots showing average module score (expression) of the top 10 computationally detected marker genes specific to each of the 13 different excitatory neuron clusters (ExN1-13). **B.** Dotplot showing the top 5 significant GO biological process terms enriched among the top 50 marker genes of the excitatory and inhibitory neuron clusters.

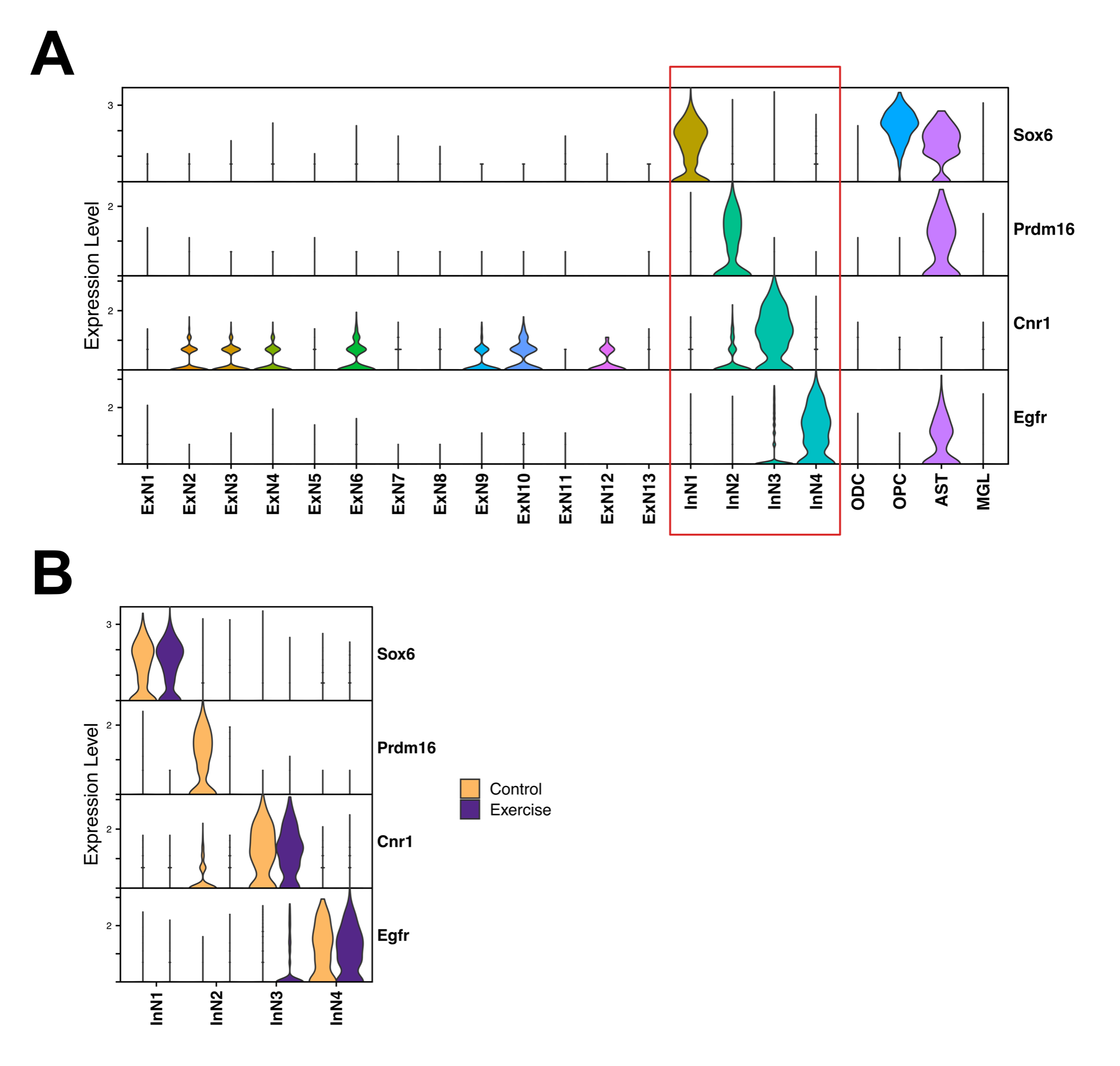

**Supp. Fig. 3: A.** Violin plots showing normalized expression of the top marker gene specific to each inhibitory neuron cluster in all cell-types *(ExN1-13: excitatory neurons; InN1-4: inhibitory neurons; ODC: oligodendrocytes; OPC: oligodendrocyte precursor cells; AST: astrocytes; MGL: microglia)*. **B.** Violin plots showing normalized expression of the top marker gene specific to each inhibitory neuron cluster (as above), in only the inhibitory clusters, further split by experimental condition.

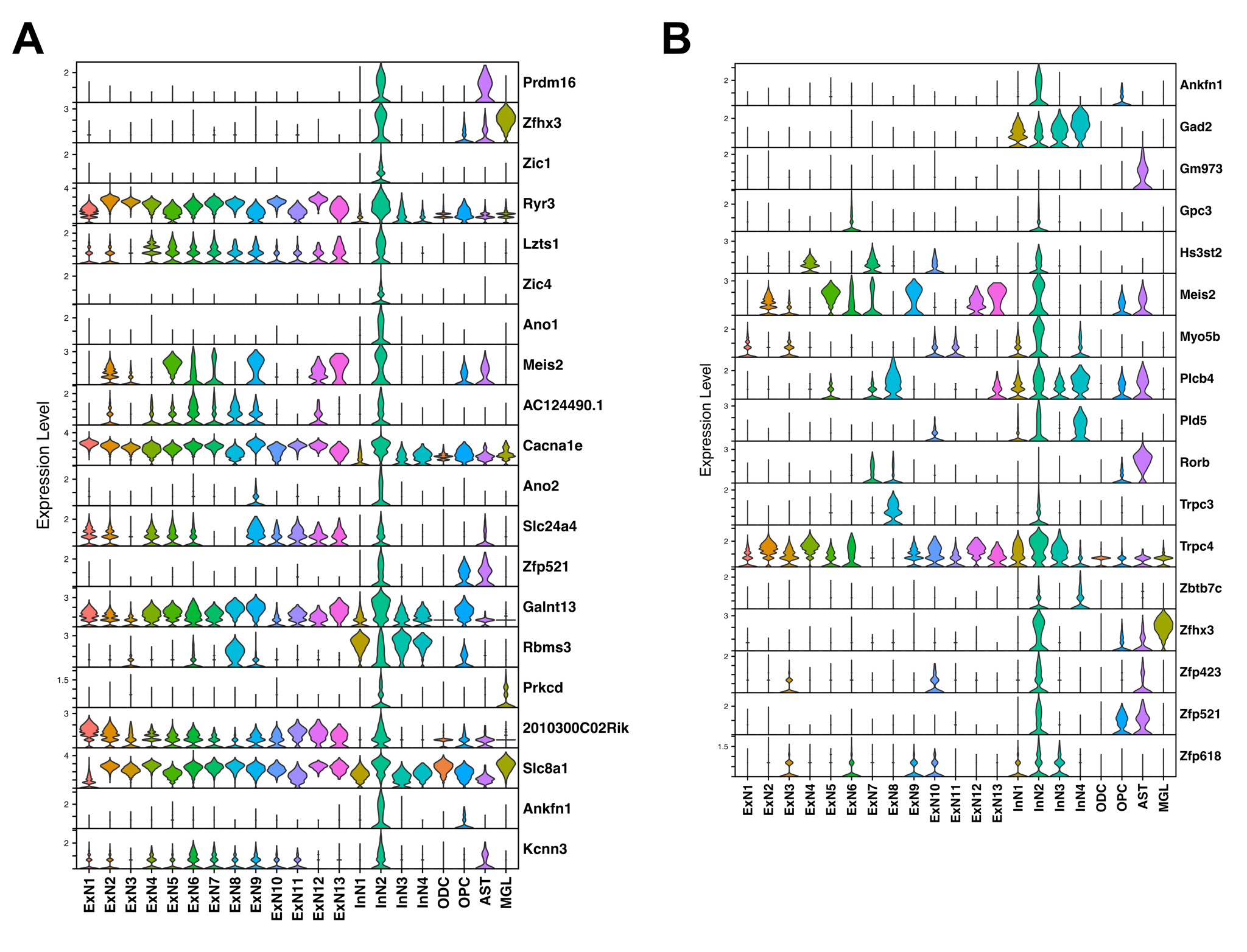

**Supp. Fig. 4: A.** Violin plots showing normalized expression of the top 20 differentially upregulated genes in the InN2 cluster, as compared to the other inhibitory neuron clusters. **B.** Violin plots showing normalized expression of the 17 genes comprising the Prdm16 regulon.

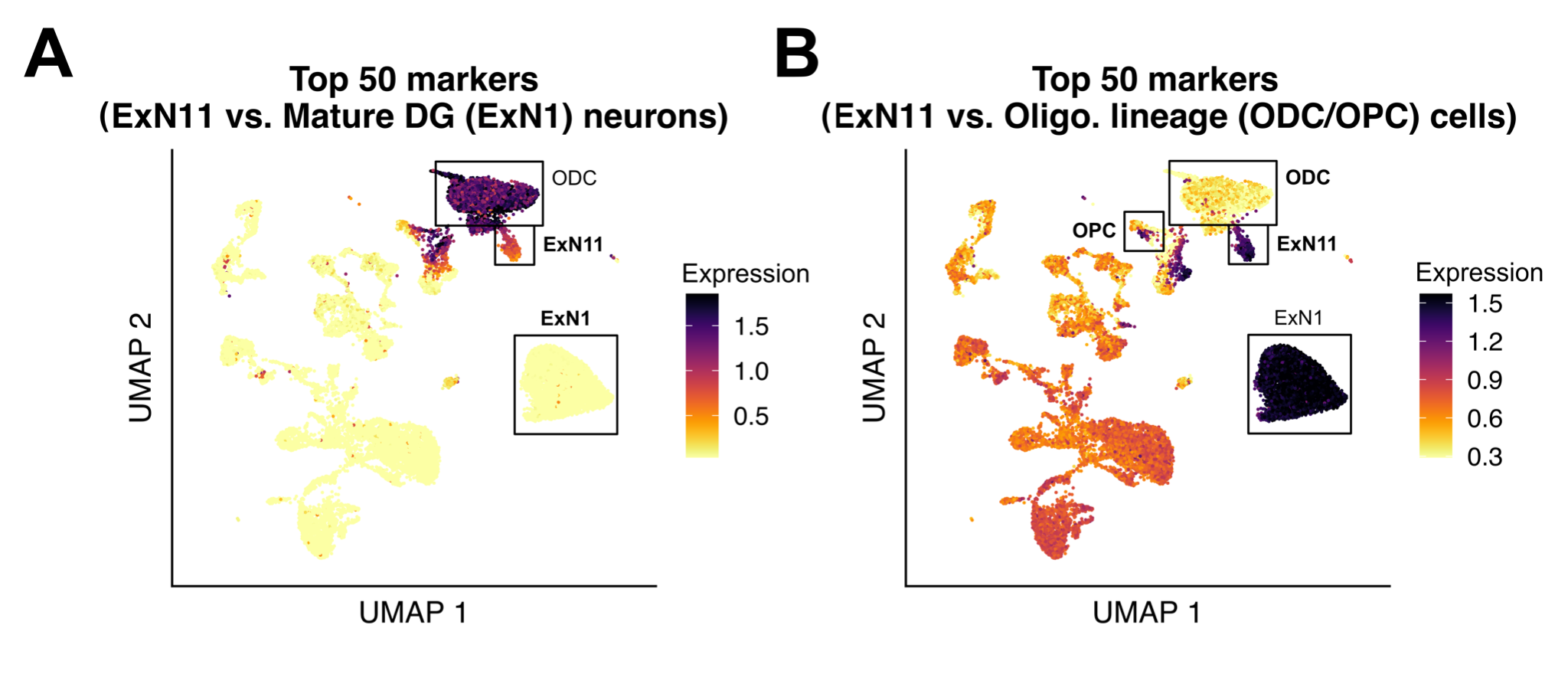

**Supp. Fig. 5: A.** UMAP plot depicting the average module score (expression) of the top 50 genes differentially expressed between the ExN11 cluster and the mature DG excitatory neuron cluster (ExN1). **B.** UMAP plot depicting the average module score (expression) of the top 50 genes differentially expressed between the ExN11 cluster and the oligodendrocyte and OPC clusters (ODC and OPC).

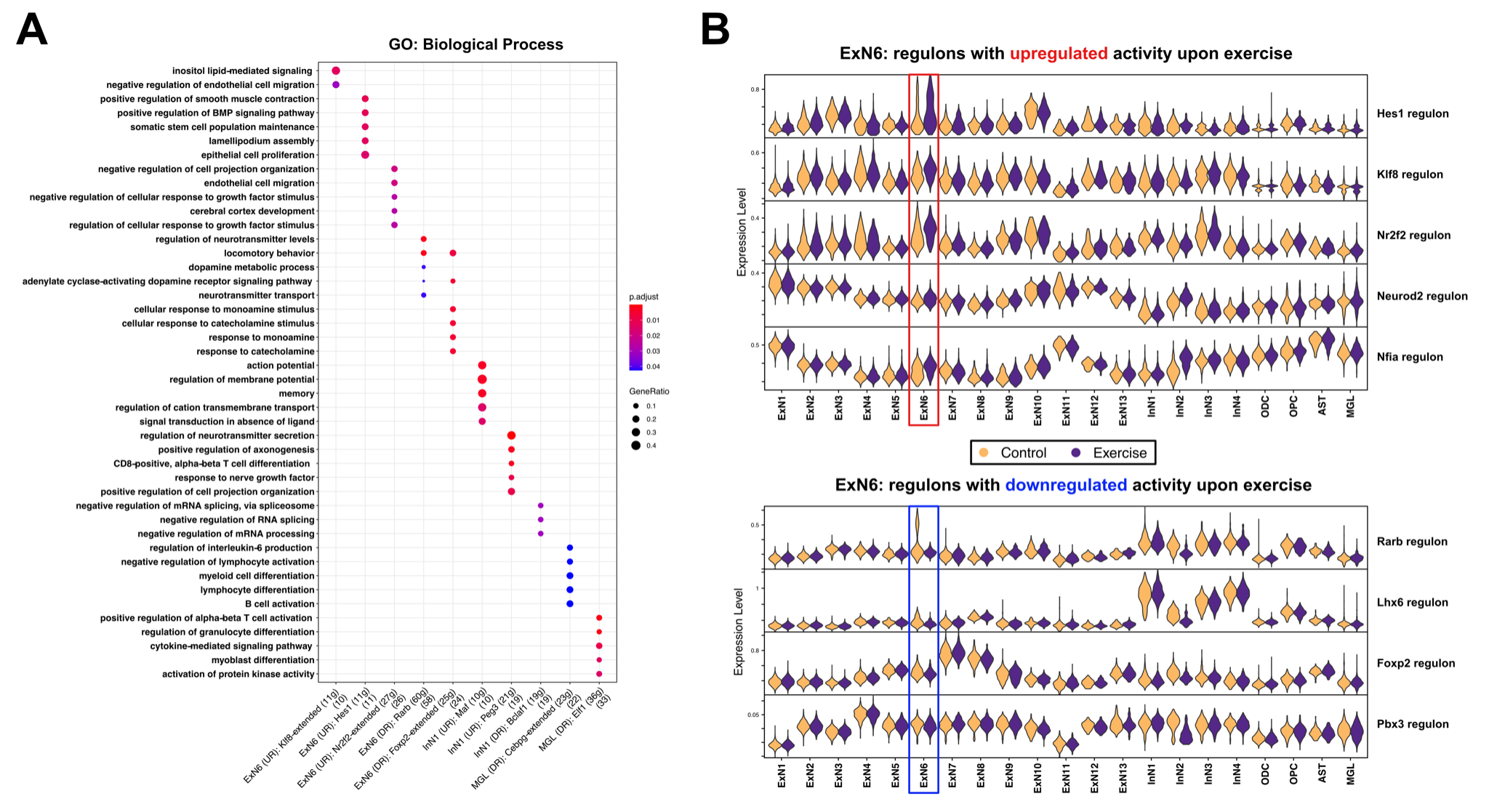

**Supp. Fig. 6: A.** Dotplot showing the top (upto 5) significant GO biological process terms enriched among genes making up the regulons with significant differential activity between cells from exercise and control conditions, in specific clusters as indicated in Fig. 4(C). **B.** Violin plots showing average module score (expression) of regulons with significant differential activity upon exercise, in the ExN6 cluster.

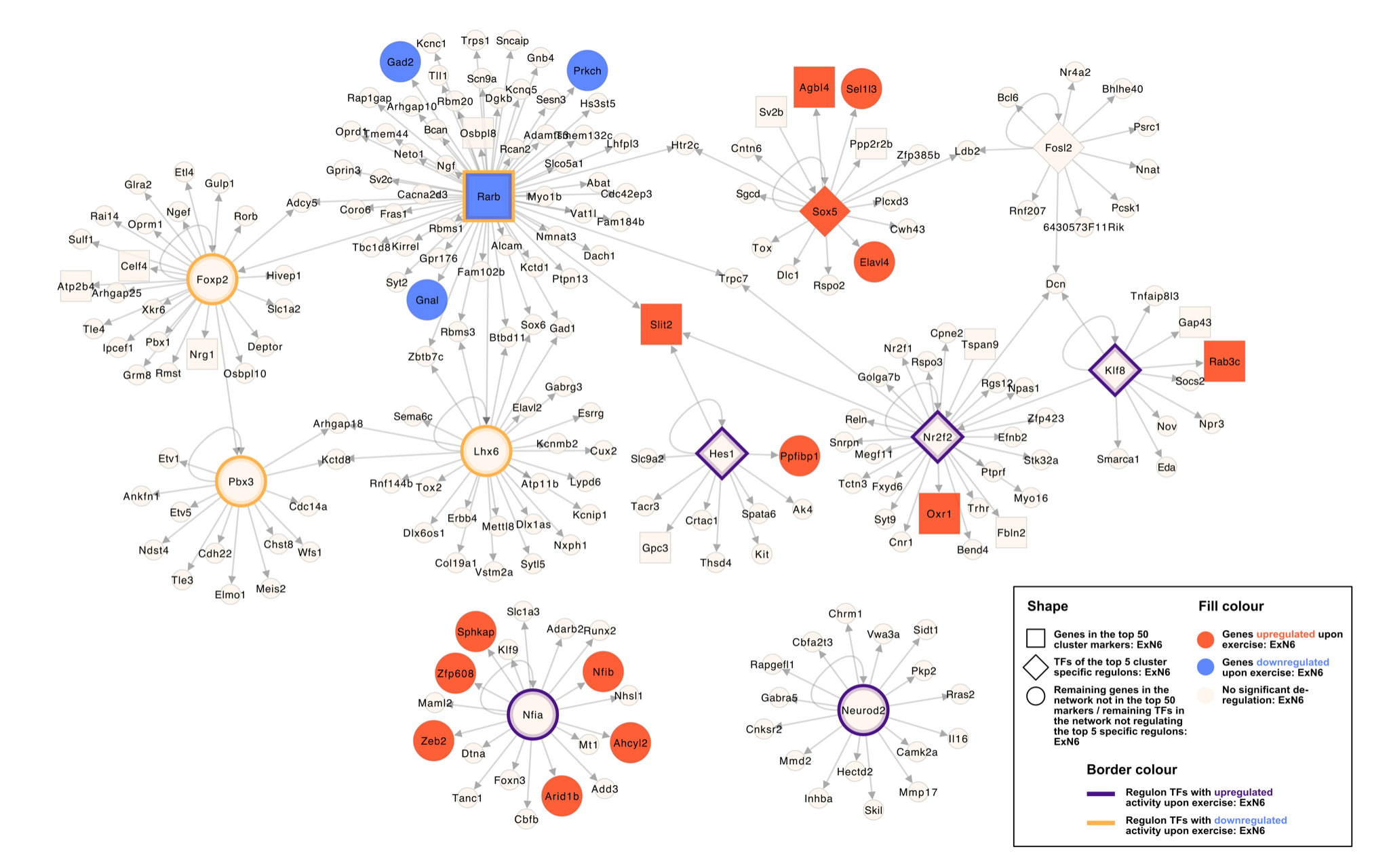

**Supp. Fig. 7:** Network plot with TFs and their respective regulon genes, specific to or showing differential activity in ExN6 upon exercise, indicating gene regulatory interactions in ExN6 cluster (Genes/TFs that are significantly deregulated upon exercise or are specifically overexpressed in ExN6 are represented bigger in size, while selected genes belonging to these regulons which are deregulated in the same direction as the regulon TF are colored in red or blue).
